# Navigation and the Efficiency of Spatial Coding: Insights from Closed-Loop Simulations

**DOI:** 10.1101/2023.01.10.523425

**Authors:** Behnam Ghazinouri, Mohammadreza Mohagheghi Nejad, Sen Cheng

## Abstract

Spatial learning is critical for survival and its underlying neuronal mechanisms have been studied extensively. These studies have revealed a wealth of information about the neural representations of space, such as place cells and boundary cells. While many studies have focused on how these representations emerge in the brain, their functional role in driving spatial learning and navigation has received much less attention. We extended an existing computational modeling tool-chain to study the functional role of spatial representations using closed-loop simulations of spatial learning. At the heart of the model agent was a spiking neural network that formed a ring attractor. This network received inputs from place and boundary cells and the location of the activity bump in this network was the output. This output determined the movement directions of the agent. We found that the navigation performance depended on the parameters of the place cell input, such as their number, the place field sizes, and peak firing rate, as well as, unsurprisingly, the size of the goal zone. The dependence on the place cell parameters could be accounted for by just a single variable, the overlap index, but this dependence was nonmonotonic. By contrast, performance scaled monotonically with the Fisher information of the place cell population. Our results therefore demonstrate that efficiently encoding spatial information is critical for navigation performance.

## 1 Introduction

Spatial navigation is critical for the survival of most animals and many neural populations in the brain appear to have evolved to support spatial learning and navigation, e.g. place cells (PC) (O’Keefe & Nadel, 1978), boundary cells (BC) Hartley, Burgess, Lever, Cacucci, and O’keefe (2000), and head direction cells (Taube, 1998). Place cells in the hippocampus are selectively active when the animal is located in one particular part of an environment, the place field (O’Keefe & Nadel, 1978). A large number of studies have examined which aspect of the environment determines the firing patterns of place cells, e.g., there is a larger number of place fields with smaller sizes close to goal radius locations as compared to elsewhere in the environment (Ainge, Tamosiunaite, Woergoetter, & Dudchenko, 2007; Dupret, O’neill, Pleydell-Bouverie, & Csicsvari, 2010; Gauthier & Tank, 2018; Grieves, Duvelle, & Dudchenko, 2018; Grieves, Wood, & Dudchenko, 2016; Hollup, Molden, Donnett, Moser, & Moser, 2001; Jarzebowski, Hay, Grewe, & Paulsen, 2022; Kaufman, Geiller, & Losonczy, 2020; I. Lee, Griffin, Zilli, Eichenbaum, & Hasselmo, 2006; J.S. Lee, Briguglio, Cohen, Romani, & Lee, 2020; Parra-Barrero, Diba, & Cheng, 2021; Sato et al., 2020; Tryon et al., 2017; Turi et al., 2019; Zaremba et al., 2017). The same observation was made near walls as compared to further away Tanni, De Cothi, and Barry (2022). Furthermore, the range of some place cell parameters can be large, e.g., place field sizes cover two orders of magnitude, ranging from tens of centimeters to more than 10 meters (Eliav et al., 2021; Kjelstrup et al., 2008; Rich, Liaw, & Lee, 2014). It is largely unknown why this variability occurs.

In sensory systems, it has been suggested that variability of the quality of sensory code might have an intrinsic source in the brain (Abbott & Dayan, 1999; Ralf & Bethge, 2010; Rieke, Warland, Van Steveninck, & Bialek, 1999). Barlow et al. (1961) suggested the efficient coding hypothesis, which has driven much research in sensory neuroscience. A sensory code is considered efficient, if it maximizes the amount of useful information that can be extracted about a sensory input while minimizing noise and redundancy, or irrelevant information, (Graham & Field, 2007; Wiskott & Sejnowski, 2002). Many aspects of the efficient coding hypothesis have been tested and confirmed in sensory systems (Barlow, 2001). Since place fields of hippocampal pyramidal neurons are similar to tuning curves in sensory neurons, the concept of efficient coding can readily be applied to spatial information, which we refer to as efficient spatial coding. One aspect of coding efficiency is how well place fields cover the environment, another is how many spikes place cells fire, and yet another is by how much the firing rate of place cells change as a function of spatial position. However, few studies have attempted to apply the efficient coding hypothesis to spatial representations, although there are notable exceptions (Finkelstein, Ulanovsky, Tsodyks, & Aljadeff, 2018; Mathis, Herz, & Stemmler, 2012; Rolls & Treves, 2011).

Efficient coding would benefit spatial navigation similar to how it benefits perceptual tasks. However, the purpose of efficient sensory codes has not been studied nearly as much and remains obscure (Simoncelli, 2003). Similarly, little is known about the exact functional role of place cells in spatial navigation and learning, in particular the role of place cell properties – such as field size, firing rate, and density – and their variations across the environment. At some point, it was hypothesized that place cells are important for path integration (Samsonovich & McNaughton, 1997), which tracks the animal’s position by integrating velocity signals. However, after the discovery of grid cells it was suggested that they might be a more likely candidate for maintaining a representation of the animal’s position via path integration (McNaughton, Battaglia, Jensen, Moser, & Moser, 2006). Nevertheless, even if that were true, path integration is only one of the computations that is required for navigation behavior. So, even if spatial representations were intimately linked to path integration, it remains to be clarified how neural path integration impacts spatial behavior.

Here we studied the function of place cells in spatial navigation and the role of place cell parameters using an artificial agent based on spiking neural networks that learned to navigate to a goal location. This requires simulating neuronal processes and the emerging spatial behaviors a closed-loop setup, in which neuronal activity is driven by the inputs from the environment, and which in turn affects actions. The action of the agent changes its state and the sensory inputs. We based our computational model on a computational modeling tool-chain (Jordan, Weidel, & Morrison, 2019) that uses the NEST simulator (Gewaltig & Diesmann, 2007) to simulate biological neural networks and Gym (Brockman et al., 2016) to model the interactions with the environment. In our simulation, the agent had to solve a spatial learning task similar to the Morris watermaze (Morris, 1981). We found that, as expected, increasing the goal size leads to a better navigation performance. By contrast, the dependence of performance on the parameters of the place cell population, i.e., cell number, field size, and peak firing rate, was less clear. However, we found a systematic relationship between performance and a new variable, the overlap index, which measures the degree of overlap between two neighboring place cells and which incorporates both place cell number and field size. However, this relationship is nonmonotonic. Finally, we found that the Fisher information, which describes how informative the place cell population is about the spatial location (Brunel & Nadal, 1998; Kloosterman, Layton, Chen, & Wilson, 2014), best accounts for navigation performance in our model.

## 2 Methods

For closed-loop simulations of spatial learning and memory we used the software framework CoBeL-spike (Closed-Loop Simulator of **Co**mplex **Be**havior and **L**earning Based on Spiking Neural Networks), which is available at https://github.com/sencheng/CoBeL-spike (Fig. 1). CoBeL-spike builds on a computational modeling tool-chain (Jordan et al., 2019) that includes the NEST simulator for networks of spiking neurons (Gewaltig & Diesmann, 2007), such as place cells (O’Keefe & Nadel, 1978), boundary cells Hartley et al. (2000), and head direction cells (Taube, 1998). The network model is based on a earlier neural network developed by Brzosko, Zannone, Schultz, Clopath, and Paulsen (2017) and described below. The interactions with the environment are modelled by OpenAI Gym (Brockman et al., 2016).

**Fig. 1.**
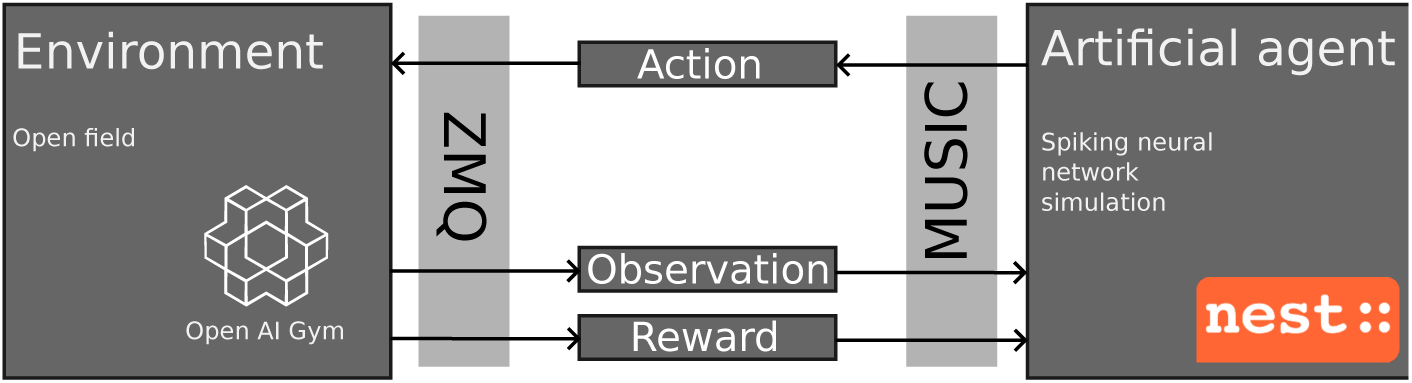
Relationship between the components of the CoBeL-spike framework. The CoBeL-spike tool-chain consists of two main components: the environment and the artificial agent. The environment is simulated using OpenAI Gym (Brockman et al., 2016) and keeps track of the agent’s position and reward and makes sure that the agent’s moves are consistent with the physical constraints. The environment sends the current state and rewards to the learning agent and receives the action from the agent. The agent is a 2-layer spiking neural network implemented using NEST (Gewaltig & Diesmann, 2007). Communication between two components is facilitated by two interfaces. MUSIC (Djurfeldt et al., 2010) transfers massive amounts of event information as well as continuous values from one parallel application to another. The action is taken through ZeroMQ (Hintjens, 2013) to the Gym environment.

### 2.1 Overview of Simulated Task

We used CoBeL-spike to simulate spatial learning in a 2.4 m *×* 2.4 m open 2-d environment (which is comparable to sizes of Morris watermazes used in experiments). Each simulation consisted of 30 trials. At the beginning of each trial, the artificial agent was located at the center of an open environment. The agent moved freely inside the environment and sought a hidden goal zone, a circular area, centered at [0.5, 0.5] from the center of the environment. Its radius *r* varied across simulations, but remained fixed within a simulation. A trial ended either if the agent entered the goal zone or after the time out of 5 s (see below), whichever occurred first. The agent was rewarded, if it entered the goal zone. This reward drove learning, so that agents tended to reach the goal zone faster in later trials. Then the next trial commenced.

The main simulation parameters that were varied independently were the goal size *r* ∈ { 0.1 m, 0.15 m, 0.2 m, 0.3 m }, the number of place cells in the input from *N_PC_* = 3^2^ to *N_PC_* = 101^2^, and the place field size in the range of *σ_PC_* = 0.02 m to *σ_PC_* = 1.4 m. Varying the last two parameters allowed us to study the role of the place cell population in spatial learning. For each parameter set, results were averaged over 100 simulations, in each of which the network was randomly initialized to simulate learning in different artificial agents. The number was chosen based on our experience that it reduced the variability of average performance to an acceptable level.

We used two variables to evaluate navigation performance of the agent in the simulated task: escape latency (*τ*) and hit rate. The former was determined by averaging the trial duration over the 100 repetitions of the same simulation. The lower *τ*, the better the navigation performance. The other performance variable was the hit rate, which was defined as the fraction of the successful trials out of the 100 repetitions of the same simulation. A trial was considered successful, if the agent found the reward within a certain time limit (see below). The higher the hit rate, the better the performance. Since both parameters measure performance, they are correlated to some extent, but that correlation can be high or low depending on the task and the behavior of the agents. For instance, if all agents learn the same trajectory to the reward, if they find it at all, then the latency and hit rate would be highly anti-correlated. This case is quite likely in simple task where there is one solution that is easily found and clearly better than other options. On the other hand, if there are multiple potential solutions that vary in their quality, it is conceivable that agents with the same success rate would show different latencies.

In preliminary simulations, we found that in most cases, the agent either found the reward within 3 s or did not find it at all. In most of the unsuccessful trials, the agent became stuck in a loop or failed to move. To make sure that simulation time was not wasted on these kinds of unsuccessful trials, we implemented a timeout after 5 s, which allows for some variation in the behavior on successful trials. If the trial duration was less than 4.5 s (there were very few data points between 4.5 s and 5 s), the trial was considered successful. Otherwise, the trial was considered unsuccessful.

### 2.2 Model Architecture

In this section, we give an overview of the components in our computational model, which are described in more detail in Fig. 2. We performed our simulations with a time step of 0.1 ms. The agent’s position **x**(*t*) modulated the firing rate 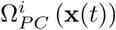 of *N_PC_* place cells (Fig. 2A), whose place field centers were evenly distributed across the environment (Fig. 2B). In addition, the input layer included eight boundary cells with firing rate 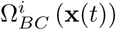 (Fig. 2C). The output spikes of place and boundary cells were fed to 40 action selection neurons (Fig. 2D). The action selection neurons formed a ring attractor network. The location of the bump in this network determined the allocentric direction of the movement at each time step. A symmetric spike-timing-dependent plasticity (STDP) enabled learning to the feedforward weights if the goal is found (Fig. 2E), mimicking the modulation of synaptic plasticity upon dopamine release (Brzosko et al., 2017) and the agent is not punished if it fails to obtain the reward. We utilized only positive reinforcement, similar to the Morris water maze paradigm (Morris, 1981).

**Fig. 2.**
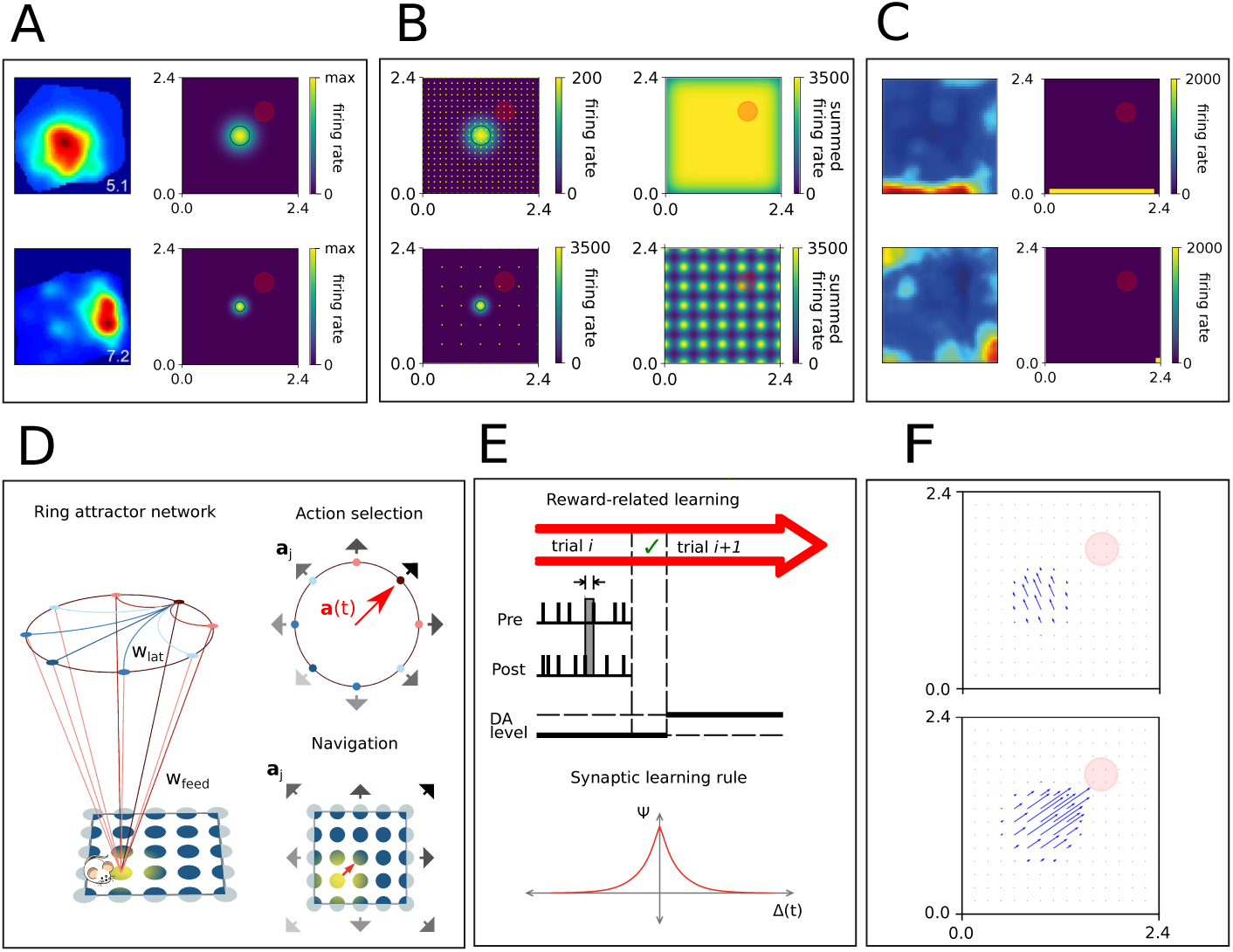
Illustration of model components. **A**: Place cells of different field sizes. Left: firing maps of two CA1 place cells (Herzog et al., 2019). Firing rate is indicated by color (from blue to red). Right: two sample firing fields of place cells in the model (top: *σ* = 0.2 m and bottom: *σ* = 0.1 m). The red disk is the goal zone (*r* = 0.2 m). Once the agent enters to this zone, the trial is terminated and the agent is rewarded as described in section 2.5. **B**: Left: Firing rate maps of two individual place fields of different sizes at the center of the environment (top: *σ* = 0.2 m, bottom: *σ* = 0.1 m). Also shown is the distribution of place field centers of the other cells (top: *N* = 441, bottom: *N* = 49, marked with yellow dots). Right: The summed firing rate map over all cells. The maximum firing rate of individual cells is chosen such that the summed firing rate at the center of the environment is 3500 Hz. **C**: Left: Firing rate maps of two boundary cells from MEC (van Wijngaarden, Babl, & Ito, 2020). Right: two boundary cells from our model. **D**: Illustration of the model architecture. Left: Place cells and boundary cells encode the position **x**(*t*) of the agent and provide input to 40 action selection neurons, which determine the direction of movement of the agent. Each action selection neuron represents one direction, which are distributed homogeneously. The action selection neurons form a ring attractor network, in which each cell excites neighboring cells (red) and inhibits globally (blue). Top-Right: In this example, the feedforward inputs are strongest to the action selection neurons that represent the North-East direction (dark red). Hence, the bump forms around that neuron and the agent moves in that direction. Bottom-Right: A different representation of the same situation that shows the preferred movement direction at different locations in the environment. **E**: Feedforward weights between place cells and action selection neurons undergo synaptic plasticity. After an unsuccessful trial the weights remain unchanged. After finding the goal, weights from place cells to action selection neurons (top) are updated according to a symmetric STDP rule (bottom). **F**: Example of the postsynaptic effect of one place cell onto action selection neurons. This impact is shown before (top) and after (bottom) learning.

### 2.3 Spatial Representations in the Input Layer

The key question in our study is what role spatial representations encoded by the place cell population play in spatial learning and navigation. The fields of the place cells in our simulations were evenly distributed across the 2-d environment on a square mesh. The spiking activity of the *i*-th place cell is modeled as an inhomogeneous Poisson process, implemented as a Poisson generator (*poisson generator*) in NEST (Gewaltig & Diesmann, 2007). Thus, the spiking dynamics of place cells was not modelled. The firing rate of each place place cell was given by

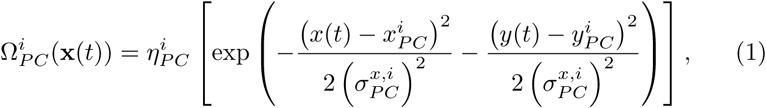

where 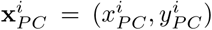 is the coordinate of the place field center of the *i*-th cell, 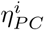 is the maximum firing rate of that cell, and 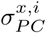 and 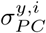 are the standard deviations of the Gaussian kernel along the *x*- and *y*-axis, respectively.

We used a simplified version of Eq. 1, where the maximum firing rate of all cells are equal and the standard deviations of the Gaussian kernels all have the same value in both directions.

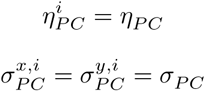

Hence, the firing map of the *t*-th place cell in our simulations can be simplified to

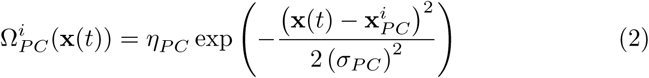

Figure 2A shows the firing rate map of two sample place cells with field sizes *σ* = 0.1 m and *σ* = 0.2 m.

Changing the place cell number *N_PC_* and field size *σ_PC_* changes the total spiking activity fed forward to the action selection neurons. This change causes a major impact on the agent’s behavior. If the input is very low, the action selection neurons do not activate the ring attractor. If the input is too high, the ring attractor is saturated and there is too much activity in the network and all action selection neurons become active. In both cases, a clear bump is not formed and the agent does not move properly. Our baseline for place cell number and place field size were *N_PC_* = 21^2^ and *σ_PC_* = 0.2 m, respectively. Preliminary simulations showed that the best performance for these parameter values occurred when the maximum firing rate of a single neuron was *η_PC_* = 200 Hz. In this case, the summed firing rate across all place cell at the center of the environment is 3500 Hz. Therefore, in all simulations we adjusted the maximum firing rate of individual cells such that the summed firing rate across all place cell at the center of the environment is 3500 Hz (see Fig. 2B), if possible. The maximum firing rate of individual cells so obtained are shown in Fig. 3.

**Fig. 3.**
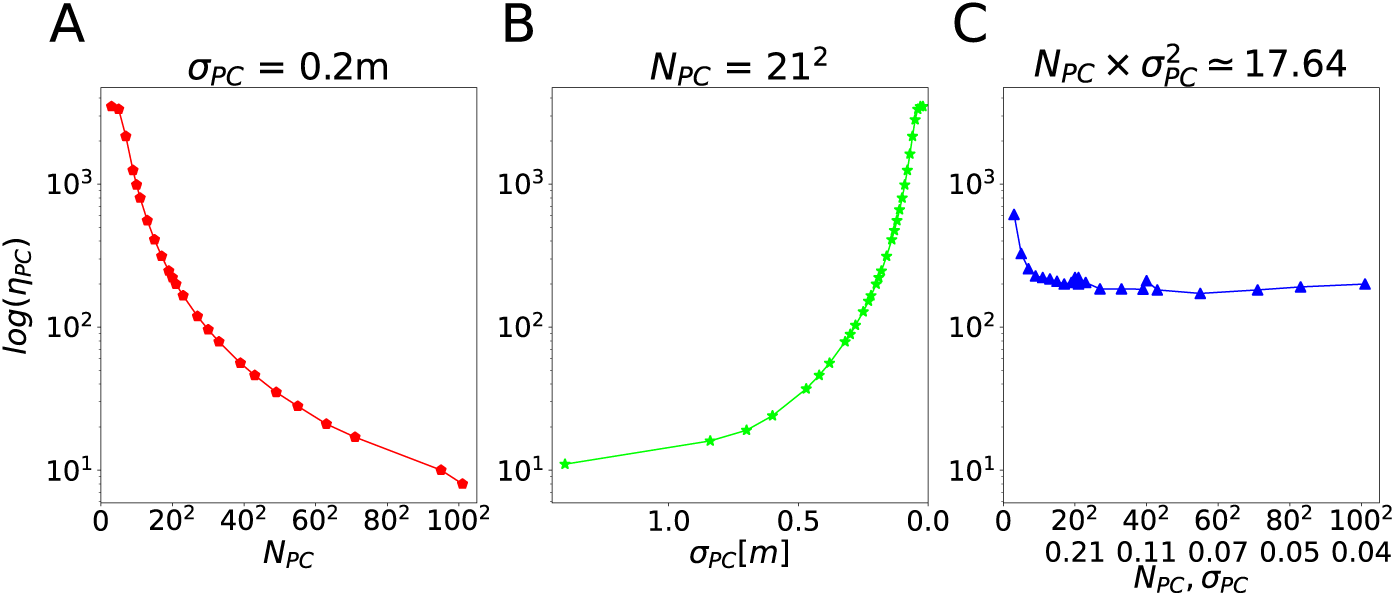
Scaling of maximum firing rate of place cells. To ensure that there was appropriate excitatory drive to the action selection network, we adjusted the maximum firing rate of place cells such that the summed firing rate at the center of the environment was 3500 Hz. Shown are the required maximum firing rates for three different simulation sets. **A**: Variable place cell number *N_PC_* for fixed *σ_PC_* = 0.2 m. We explore such different peak firing rates (8 *∼* 3500 Hz), because the firing rates of all fields at the center change when *σ* is adjusted. For higher values of *σ*, more fields contribute to the summed firing rate at the center. **B**: Variable place field size *σ_PC_* for fixed *N_PC_* = 21^2^. **C**: Both cell number and field size change simultaneously, such that *N_PC_ × σ*^2^ = 17.46.

At first glance, these maximum firing rates might appear unrealistically high. However, for computational reasons our simplified model includes far fewer neurons than the hippocampus of rodents. We regard each neuron in our simulation as representative of a larger number of biological place cells that all have a place field in the same location. A rough back-of-the-envelop calculation shows that the total number of spikes that all place cells send to action selection neurons at a given moment in our model (3500 Hz) roughly matches the spike rate that downstream neurons might receive from hippocampal neurons. We assume that a neurons received projections from about 10,000 place cells, of which about 25% are active in a typical experimental environment. Assuming further that each place field covers about 15% of the environment’s area yields about 375 neurons that are simultaneously active. If each neuron fires with a maximum firing rate of 10 Hz, the total input rate is about 3750 Hz.

To make sure that the agent does not get stuck at the borders of the environment, we added eight boundary cells that allowed us to model the obstacle avoidance behavior of animals. Boundary cells were also modelled as inhomogeneous Poisson processes. The firing rate map of each boundary cell was rectangular of size 2*x_hw_ ×* 2*y_hw_* (*hw* stands for half width) centered at 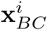:

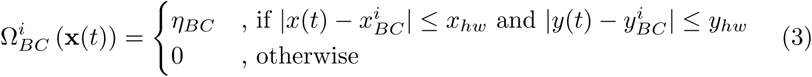

The centers of four boundary cells are located in the middle of each border and the centers of four further border cells are located at each corner. The firing field maps of two sample boundary cells are shown in Fig. 2C. The boundary cells had a fixed projection pattern to the action selection neurons, such that the agent was repelled from the border or corner towards the inside of the environment.

### 2.4 Decision Making With Action Selection Neurons

At the core of the model network were 40 action selection neurons that formed a ring attractor. Each action selection neuron represents one preferred direction of movement *θ^j^*, distributed homogeneously over 360°. The cells were connected to each other though lateral synaptic weights such that neurons with similar preferred directions excite each other and neurons coding different movement directions inhibit each other (Brzosko et al., 2017), i.e.,

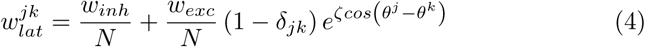

where *w_inh_* = *−*400 and *w_exc_* = 50 are inhibitory and excitatory weights, respectively, *δ* is the Kronecker delta, and *ζ* = 20. The lateral weights remained fixed throughout all simulations.

The parameter values ensured that the network is a continuous attractor network, where the attractor is an activity bump. That is, at any given point in time only a subset of neurons with similar movement directions was active and the rest of neurons were silent due to strong inhibition from the active neurons.

Feedforward inputs project from the place and boundary cells to action selection neurons. At the beginning of each simulation, the feedforward weights 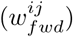 between input neurons and action selection neurons were randomly drawn from a normal distribution with mean 30 and standard deviation 5. The feedforward weights were plastic, and could be potentiated if the agent found the goal, up to a maximum of 60 (see below).

To visualize the postsynaptic effect of place cells and boundary cells onto action selection neurons, we represented each action selection neuron by a unit vector that points in the preferred direction of that neuron (**a***^j^*) and then computed the sum over all such unit vectors weighted by the synaptic input that the neurons received from the input layer in a particular location (Fig. 2F).

We modelled neurons of the action selection network as leaky-integrate-and-fire’ (LIF) neurons. The dynamics of sub-threshold membrane potential (*v^i^*) of the *j*-th LIF neurons is (Spreizer, Aertsen, & Kumar, 2019):

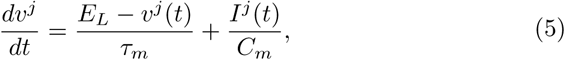

where 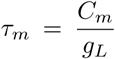 denotes the membrane time constant, *C_m_* the membrane capacitance, *g_L_* the leak conductance, *E_L_* the leak reversal potential, and *I^j^*(*t*) the total synaptic input current. The input current is the convolution of spikes and an alpha function (Bernard, Ge, Stockley, Willis, & Wheal, 1994; Jack, Noble, & Tsien, 1983; Rotter & Diesmann, 1999). The LIF neurons were implemented using the *iaf psc alpha* neuron in NEST (Diesmann, Gewaltig, Rotter, & Aertsen, 2001; Gewaltig & Diesmann, 2007; Kobayashi, Tsubo, & Shinomoto, 2009; Rotter & Diesmann, 1999; Yamauchi, Kim, & Shinomoto, 2011). The values of the model parameters are summarized in Table 1.

**Table 1.** Parameters of LIF neurons, implemented for the action selection neurons

| parameter | value | unit | description |
| --- | --- | --- | --- |
| E<sub>L</sub> | -70 | mV | Resting membrane potential |
| C<sub>m</sub> | 250.0 | pF | Capacity of the membrane |
| tau<sub>m</sub> | 10.0 | ms | Membrane time constant |
| t<sub>ref</sub> | 2.0 | ms | Duration of refractory period |
| V<sub>th</sub> | -55 | mV | Spike threshold |
| V<sub>reset</sub> | -70.0 | mV | Reset potential of the membrane |
| tau<sub>syn_ex</sub> | 5 | ms | Rise time of the excitatory synaptic alpha function |
| tau<sub>syn_in</sub> | 5 | ms | Rise time of the inhibitory synaptic alpha function |
| I<sub>e</sub> | 0.0 | pA | Constant input current |
| V<sub>min</sub> | -inf | mV | Absolute lower value for the membrane potential |

To translate the activity of action selection neurons to navigation, the unit vectors **a***^j^* representing the action neurons’ preferred direction were weighted by the exponentially discounted spikes times and summed to obtain the movement vector selected by the network **a**(*t*) (Brzosko et al., 2017).

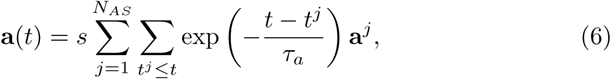

where *s* = 10*^−^*^4^ m is the step size per spike and *N_AS_* = 40 the number of action selection neurons, and *τ_a_* = 0.5 ms. The second sum runs over all spikes that the action selection neuron *j* fired before the current time *t*. This scheme made it possible for the agent to move freely in any direction with different step sizes.

The activity of the action selection neurons was not changed between trials, i.e., the activity of the action selection neurons in the last moment of trial *t* is the same as at the beginning of the next trial *t* + 1. Therefore, the initial conditions of trials are different from one another. This made it possible for the agent to find the goal after an unsuccessful trial, because otherwise the agent would repeat the same behavior on the next trial since weights do not change in/after unsuccessful trials.

### 2.5 Synaptic Plasticity Rules

The feedforward weights from place and boundary cells to action selection neurons play a critical role in our model. They define the policy of the agent. Plasticity of these weights enables the agent to learn the location of the goal zone. We applied a modified STDP rule that uses an all-to-all spike pairing scheme and eligibility trace that allows for delayed update. The following learning rule was applied to feedforward weights, if the agent finds the goal (Izhikevich, 2007; Potjans, Morrison, & Diesmann, 2010).

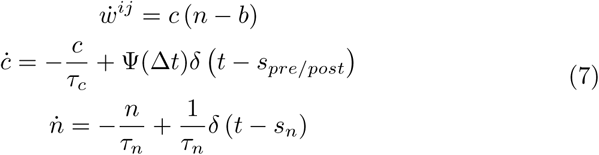

where *ẇ* denotes the change applied to the feedforward weight connecting the *i*-th place cell to the *j*-th action selection neuron. *c* is an eligibility trace that allows for delayed updates, which is necessary and sufficient to learn causal associations between synaptic activity when outcomes are delayed. This concept is a fundamental component of reinforcement learning network models (Brzosko et al., 2017; Florian, 2007; Izhikevich, 2007; Legenstein, Pecevski, & Maass, 2008). *n* is the dopamine concentration, and *b* is the dopamine baseline concentration. *τ_c_* and *τ_n_* are the time constants of the eligibility trace and the dopamine concentration, respectively. *δ*(*t*) is the Dirac delta function, *s_pre/post_* the time of a pre- or post-synaptic spike, *s_n_* the time of a dopamine spike.

The spiking relationship between pre- and post-synaptic neurons modulated plasticity via the following window function of additive STDP:

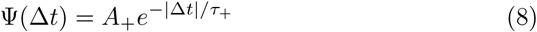

where Δ*t* = *s_post_ − s_pre_* is the temporal difference between a post- and a presynaptic spike, *A*_+_ the amplitude of the weight change, and *τ*_+_ a time constant. We used the *stdp dopamine synapse* in NEST (Gewaltig & Diesmann, 2007) to implement this model with the parameters given in Table 2.

**Table 2.** STDP learning parameters, implemented to feed forward weights from place cells to action selection neurons

| parameter | value | unit | description |
| --- | --- | --- | --- |
| A_plus | 0.002 | real | Amplitude of weight change for facilitation |
| tau_plus | 20.0 | ms | STDP time constant for facilitation |
| tau_c | 200.0 | ms | Time constant of eligibility trace |
| tau_n | 0.1 | ms | Time constant of dopaminergic trace |
| b | 0.0 | real | Dopaminergic baseline concentration |
| Wmin | 0.0 | real | Minimal synaptic weight |
| Wmax | 60.0 | real | Maximal synaptic weight |

### 2.6 Characterizing Spatial Coding in the Place Cell Population

Changing the number of place cells (*N_PC_*) and/or their field size (*σ_PC_*) changes the spatial coding in the place cell population. To quantify spatial coding we use two different measures. The first such measure, the *overlap index*, is defined by comparing two neighboring place fields, assuming that all fields are equidistant across the environment.

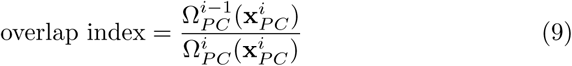

So, the overlap index is the average firing rate of one cell 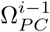 at the center of the next closest place field 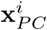, normalized by the peak firing rate of the other neuron 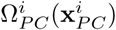. Note that by definition, the overlap index is independent of the peak firing rate of the individual neurons in our network. We also quantified spatial coding by calculating the Fisher information matrix of the place cell population (Brunel & Nadal, 1998; Fisher, 1922):

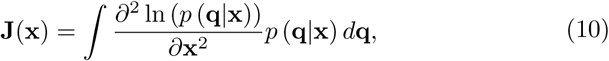

where **q** is the population activity vector. Given the place cell model, the Fisher information matrix at a location **x** is given by the following equation.

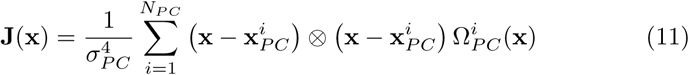

Since we use isotropic place fields, we average over the diagonal elements of the Fisher information matrix to quantify spatial coding by the place cell population. In the following, we refer to this average as the Fisher information of the place cell population.

## 3 Results

To study the functional role of the properties of place cells in spatial navigation, we performed closed-loop simulations of spatial learning controlled by a spiking neural network using the CoBeL-spike software framework. The behavior was similar to that of rodents in the Morris watermaze. Synaptic plasticity in the network based on STDP and an eligibility trace gradually changed the behavior of the agent such that it found the goal more often and faster in later trials (Fig. 4). In other words, the agent successfully learned the spatial navigation task in our computational model. The outcome of learning is also evident in more direct trajectories to the goal and the policy encoded in the feedforward projections between spatial inputs and the action selection network (Fig. 5). In the following, we study systematically how task and place cell parameters affect navigation performance on this spatial learning task.

**Fig. 4.**
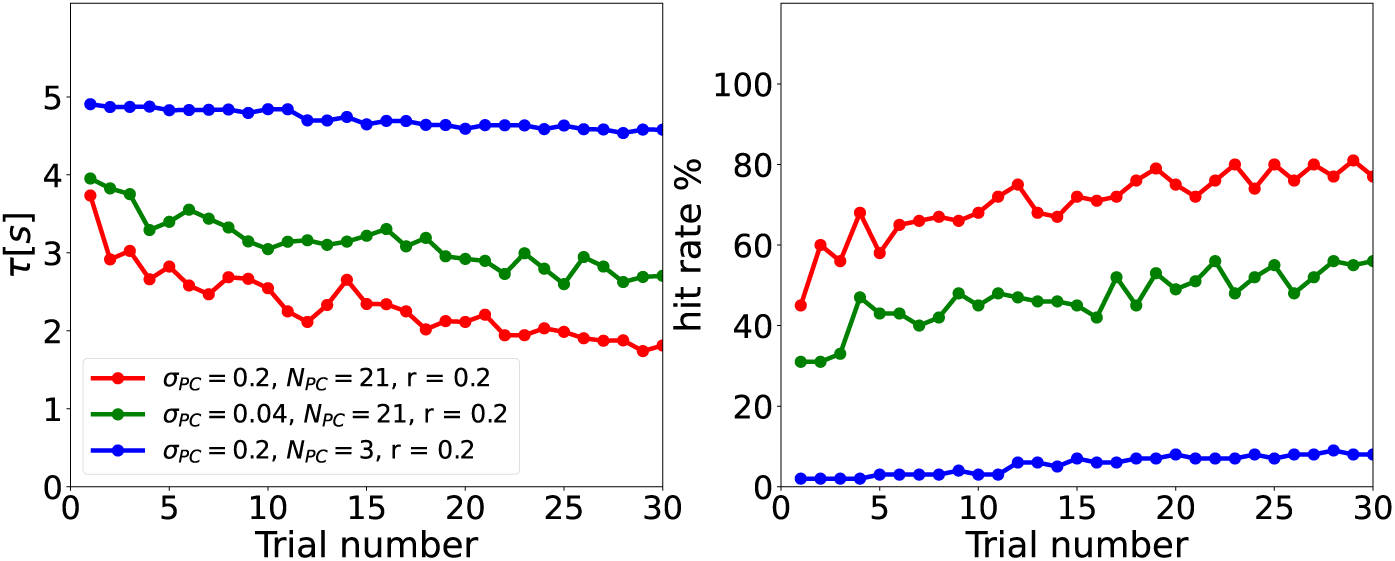
Learning curves in the simulated task are sensitive to network and task parameters. Examples of escape latencies (left column) and hit rates (right column) as a function of trial number for three sets of network and task parameters. For some parameter learning is evident in decreasing latency and increasing hit rate (red and green lines), whereas for others there is little to now learning (blue line).

**Fig. 5.**
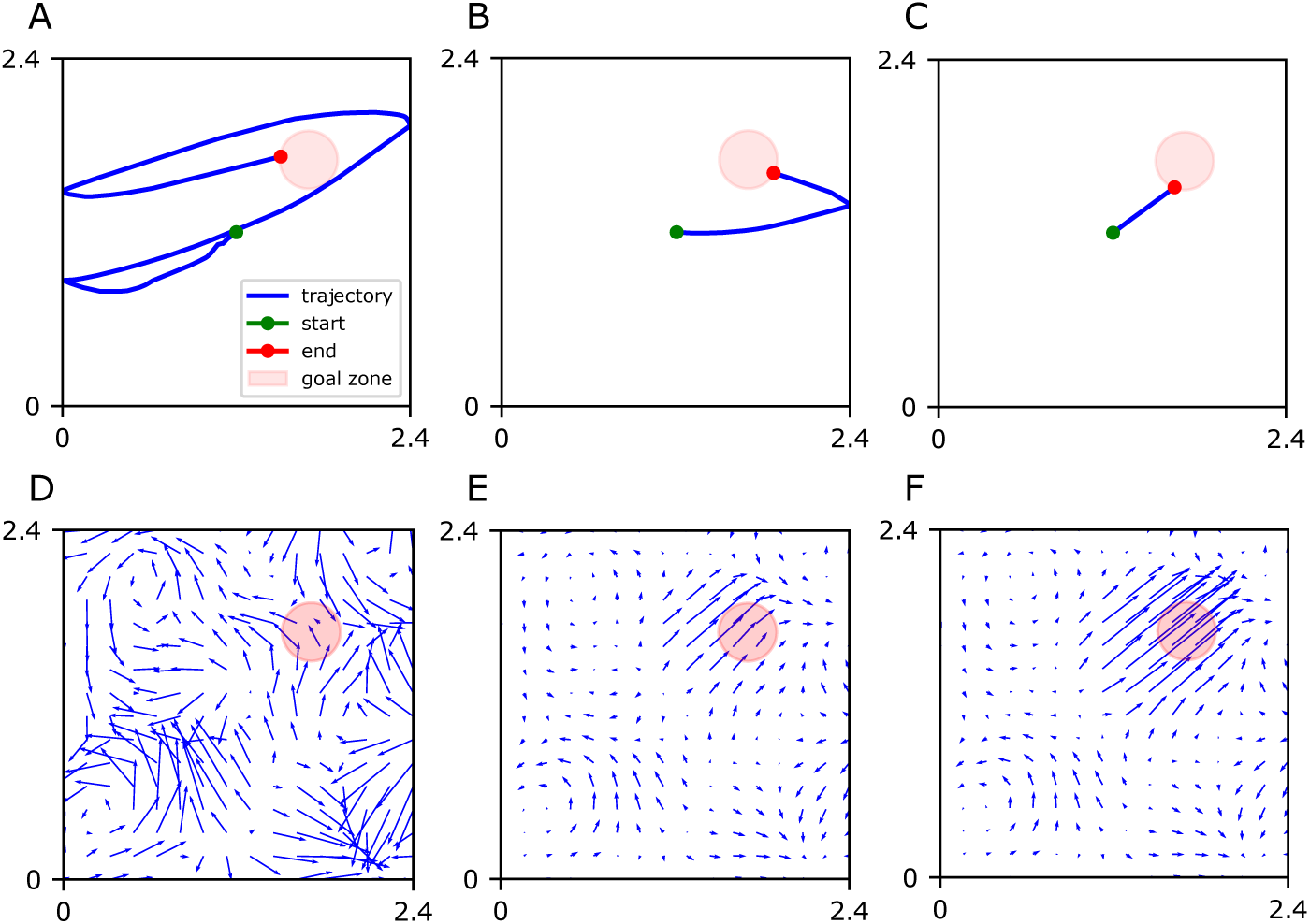
Example of behavior and learned policy during training. In the simulated task, the agent starts every trial in the center (green dot) and has to navigate to the goal zone (red disk) to receive reward. **A-C:** Sample trajectories of the agent in the first (A), fifth (B), and tenth (C) trial. **D-F:** Sum of the effects of all place cells on action selection neurons in the first (D), fifth (E), and tenth (F) trial. Arrows indicate the preferred direction of movement at each spatial location, which represents what is called the policy of the agent in reinforcement learning. Through learning, the trajectory becomes shorter and the agent finds the reward faster.

### 3.1 Studying the Impact of Cell Number, Field Size, and Goal Size on Navigation Performance

First, we kept the field size of the place cells unchanged (*σ_PC_* = 0.2 m), while we varied the place cell number from 3^2^ to 101^2^) and goal size. We adopted the place field size *σ_PC_* = 0.2 m. To ensure that the agent is initially located at the center of a place cell at the start of each trial, we consistently assigned an odd number to the place cell numbers in a row. This approach helps to drive the agent initially and maintains consistency across trials. The smallest practical number for place cell is therefore 3^2^. Our performance measures are escape latency (the lower, the better) and the hit rate (the higher, the better). We observed that the simulation results based on escape latency (Fig. 6A) and the hit rate (Fig. 6D) for this parameter set are consistent with one another. As mentioned above, this indicates that all agents that are successful find similar trajectories to the goal. Hence, we will only present the escape latency for any further computations or analyses. A clear result that we found throughout all simulations was that the agent performs better as the goal becomes larger. This occurs because the task becomes easier in two ways: the distance between start and goal becomes shorter and a less precise policy suffices to reach the goal zone.

**Fig. 6.**
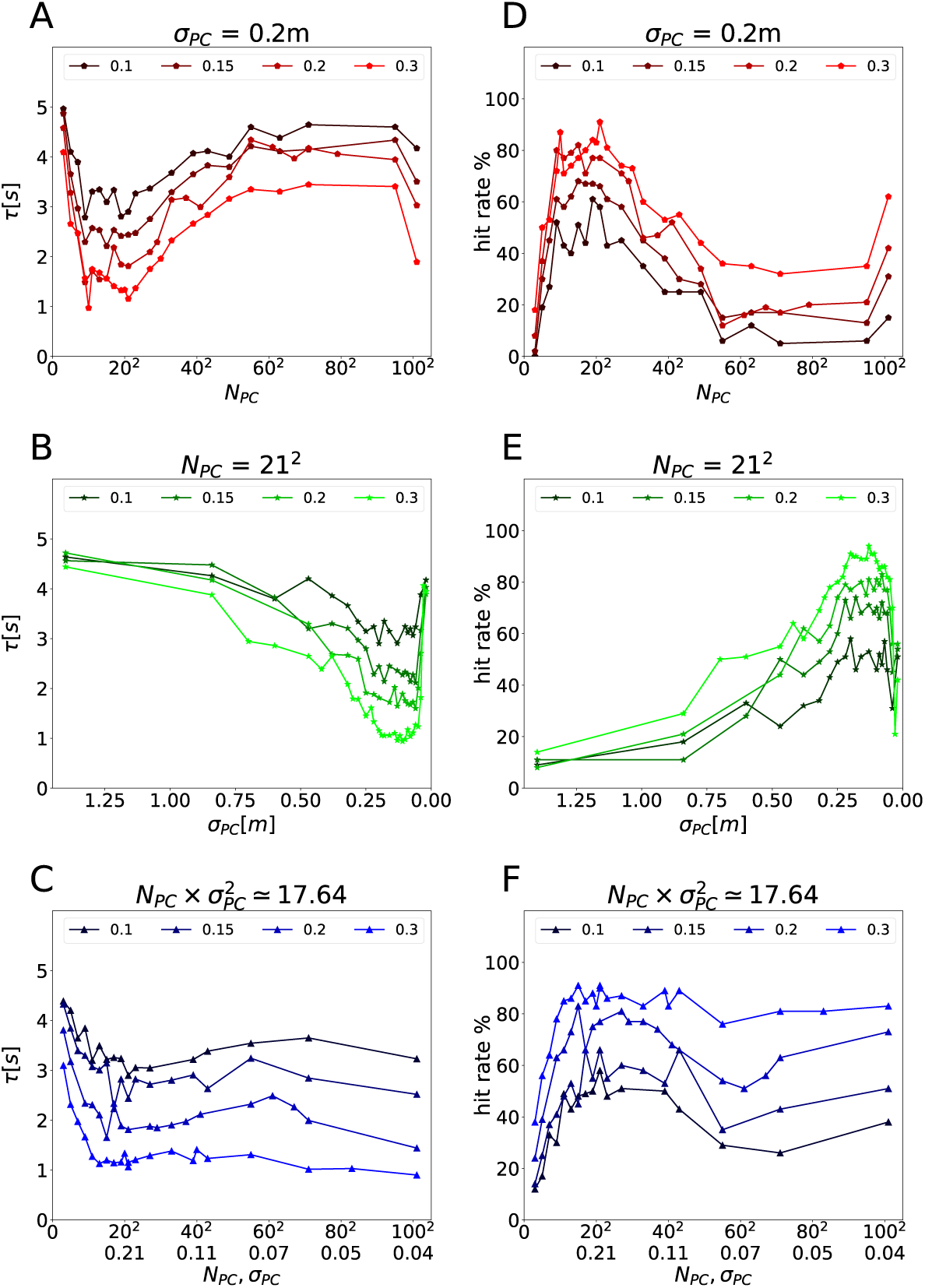
Asymptotic navigation performance after learning. **A-C**: Escape latency as a function of place cell number *N_PC_* for fixed *σ_PC_* = 0.2 (A), of place field size *σ_PC_* for fixed *N_PC_* = 21^2^ (B), and simultaneous scaling of both variables such that 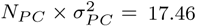 (C). Each point represents the average trial durations over 100 repetitions. The color represents different goal radii (as indicated). **D-F**: The hit rate, i.e. the fraction of trials in which the agent successfully navigates to the goal, shown using the same plotting convention as in A-C.

By contrast, the relationship between performance and cell number was less clear. Increasing the cell number increases the density of place field, which in turn improves the spatial coverage by the place cell population. So, larger cell number should result in better navigation performance. We varied the cell number from 3^2^ to 101^2^. We report cell number using this convention, because the cells are always located on a 2-d grid (Fig. 2B). The agent performed the worst for very small cell number and gradually performed better when cell number increased – as predicted (Fig.6 A,D). However, for a large number of cells, the performance declined. We hypothesize that this occurs, because the place fields have too much redundancy, i.e. overlap.

Next, we changed the place field size while keeping the cell number constant at *N_PC_* = 21^2^ (Fig. 6B,E). We chose that value because the agent performed well in the previous simulation. We expected that the smaller the fields, the more precise the spatial coding and the better the behavioral performance will be. Hence, we reversed the x-axis for this plot from the standard convention (left: *σ_PC_* = 1.4 m, right:*σ_PC_* = 0.02 m). As expected, performance was better (lower *τ* and higher hit rate) for smaller place field sizes, but, like above, there was an optimal field size beyond which the performance declines. We hypothesize that very small fields cannot properly cover the environment, which leads to issues with navigation and/or spatial learning.

### 3.2 Investigating Possible Invariance to Spatial Coverage

Taken the results of the two simulation sets together, we hypothesize that the coverage of the environment by the place cell population might be the variable that determines navigation performance. We therefore defined the *coverage index* as follow:

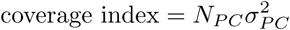

To test our hypothesis, we next changed the cell number and field size in such a way that leaves the coverage index constant at 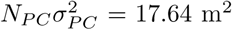. This value arises from our standard parameters for *N_PC_* = 21^2^ and *σ* = 0.2 m. Performance is roughly constant for a sizable range of place cell parameters (Fig. 6C,F), but at lower cell numbers and larger field sizes, the performance deviates quite dramatically. So, coverage alone is not sufficient to explain the difference in navigation performance.

### 3.3 Accounting for Navigation Performance Using Measures of Spatial Coding by the Place Cell Population

We hypothesized further that it is not just the spatial coverage by place cells *per se*, but more subtle properties of spatial coding by the place cell population that matter for spatial navigation. We therefore defined the overlap index to measure spatial coding by quantifying how close the peaks of neighboring place fields are relative to their width. The overlap index increases if fields move closer together while their sizes *σ_PC_* remain constant. This occurs when only the cell number *N_PC_* is increased (Fig. A1, from left to right). The overlap index also increases if field sizes increase while the field density stays constant, i.e., the cell number is constant (Fig. A1, from bottom to top).

We calculated the overlap index for the three sets of simulations that we performed above (Fig. 7, left column). Per definition, the overlap index increases from 0 to 1 with cell number (Fig. 7A), and decreases from 1 to 0 with decreasing field size (Fig. 7B). Also, as intended the overlap index deviates from the coverage index, i.e., when the coverage index is held constant while place cell parameters change, the overlap index varies (Fig. 7C)

**Fig. 7.**
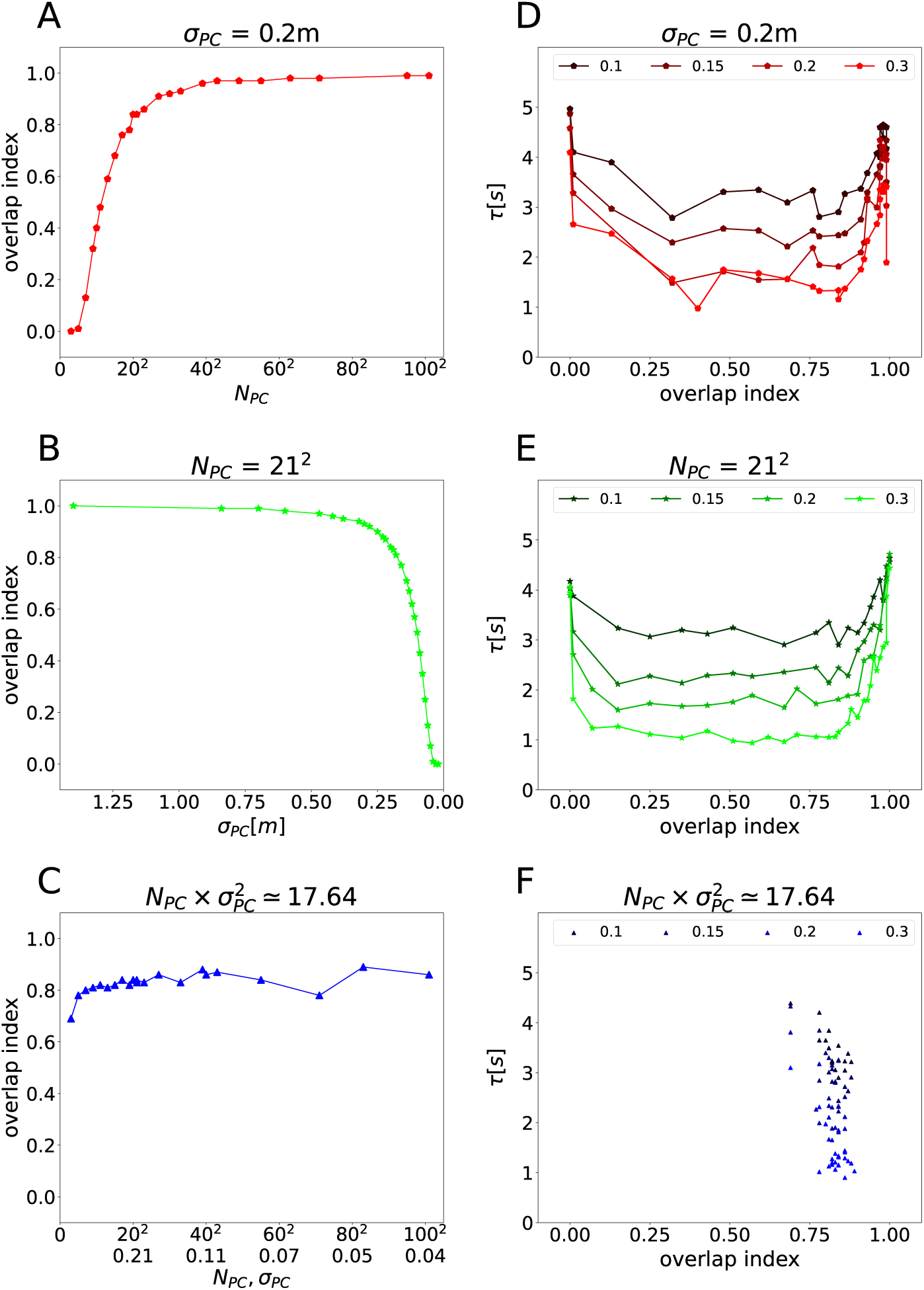
The agent’s navigation performance shows a dependence on overlap index. The dependence of the overlap index on key simulation parameters: **A**: cell number when sizes are fixed, **B**: field size when numbers are fixed, and **C**: when cell number and field sizes scale inversely to one another. **D-F:** Escape latency depends on overlap index for the three types of scaling, but the relationship is non-monotonic.

Interestingly, when we plotted navigation performance vs. the overlap index, a pattern emerged that is similar across the three simulation sets (Fig. 7, right column). However, the relationship is nonmonotomic (Fig. 7D, E). What is more, the asymmetric u-shape has a very wide bottom, so that the best performance is observed around a large range of values of the overlap index between 0.2-0.8 (Fig. 7D,E). These results are clear indications that the overlap index accounts for spatial coding features that are important for spatial navigation, but does not capture it well enough.

Next, we used a measure of sensory coding that is often used in sensory neuroscience (Brunel & Nadal, 1998): the Fisher information. We calculated the Fischer information for the place cell population at all locations between the start point (center of the environment) and the center of the reward zone (see section 2.6) and use the minimum value encountered to characterize the place coding by the population – more on this point below. The Fisher information increases if the number of cells increase while field sizes remain constant (Fig. A2, from first to second column). Since the Fisher information also takes into account the peak firing rate, we held the peak firing rate constant for this comparison. However, if we scale the peak firing rate as we do in our closed-loop simulations, the simultaneous scaling of cell number and peak firing rate leaves the Fisher information virtually unchanged (Fig. A2, from first to third column). When field sizes increase and peak firing rates simultaneously decrease, Fisher information decreases (Fig. A2, from bottom to top).

These properties of the Fisher information explain its scaling in the three simulation sets (Fig. 8, left column). When cell number is increased, and peak firing rate is decreased, while field sizes remain constant, the Fisher information remains constant (Fig. 8A). In the other two cases, the Fisher information increases as expected (Fig. 8B,C). However, for very low cell numbers or very small field sizes this trend breaks, because in these cases the place fields are so sparse that the Fisher information varies substantially between locations where there is a place field and those where there is none (see Fig. A3). This is the reason why we chose the minimum Fisher information along the path to quantify spatial coding in the place cell population.

**Fig. 8.**
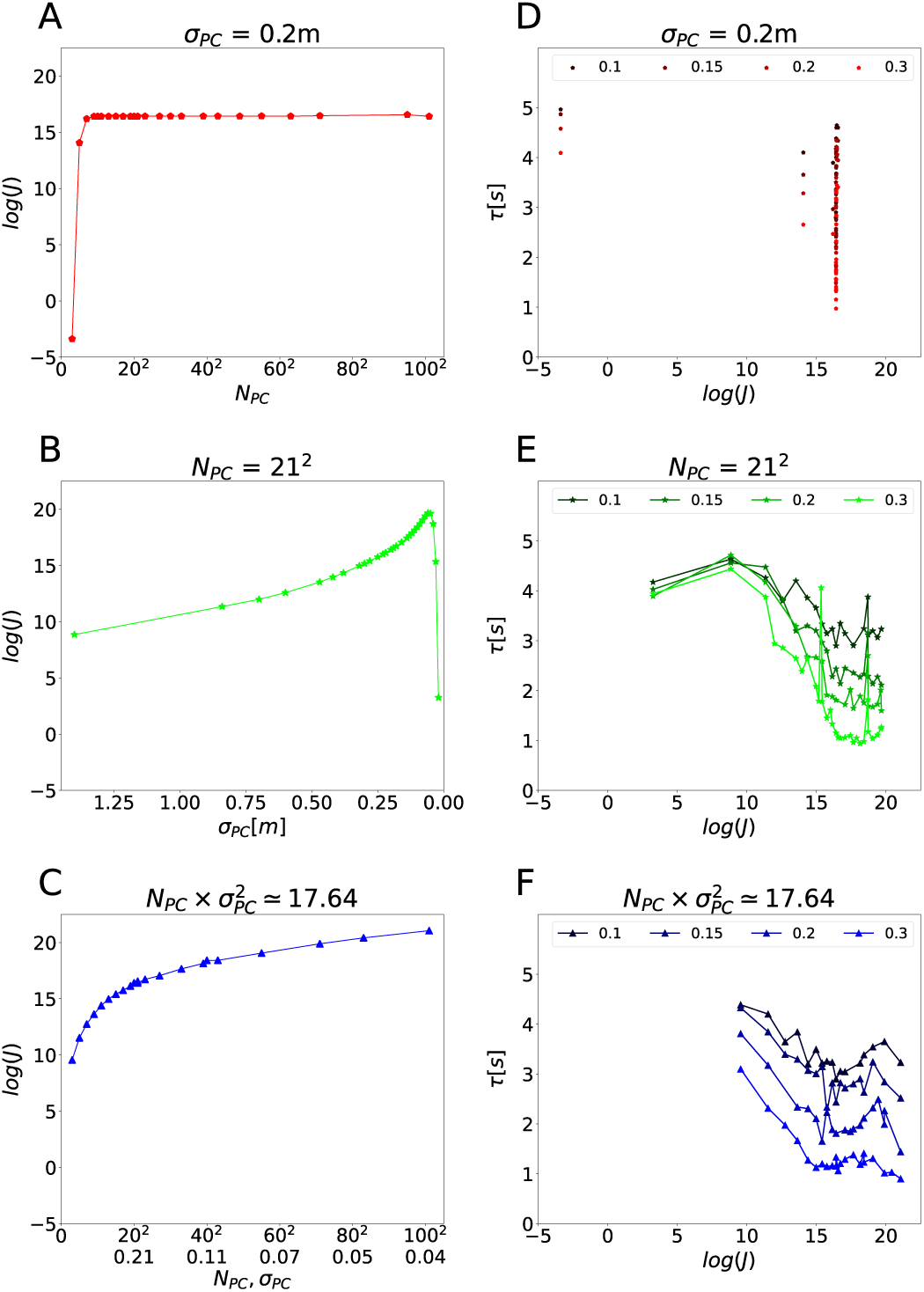
Fisher information accounts for agent’s navigation performance. Same plotting convention as in Fig. 7 for the binary logarithm of the minimum value Fisher information on the direct path between the start point and the goal’s center. **A:** Except for very low cell numbers, the Fisher information remains constant when increasing place cell number in our simulations, because we simultaneously decrease the maximum firing rate of individual cells (see Fig. A2). **B**: Decreasing the place field size lead to an increase in Fisher information, because of the power of 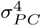 in the denominator of Fisher information. However, for extremely small place fields the Fisher information drops because the place representation becomes too sparse. **C**: Scaling of cell number and field size simultaneously also increases Fisher information. **D-F**: For the three types of parameter scaling, the escape latency monotonically decreases, and therefore the performance increases, with increasing Fisher information.

The navigation performance vs. Fisher information plots are similar across all three simulation sets (Fig. 8, right column) and, the relationship is largely monotonic, unlike the relationship between performance and overlap index. We therefore conclude that the spatial coding of the place cell population, as measured by the Fisher information, is indeed an important driving factor in the behavioral performance of the agent in the closed-loop simulations. Modifying the simulation parameters to increase the Fisher information leads to improved performance of the agent.

### 3.4 Consistent Relationship Between Navigation Performance and Efficiency of Spatial Coding

To investigate whether the relationship between navigation performance and spatial coding is the same in all three simulation sets, we plotted the results of all simulations in one figure. Both relationships navigation performance vs. overlap index (Fig. 9A) and navigation performance vs. Fisher information (Fig. 9B) are consistent across all three simulation sets. Our results therefore show that the efficiency of spatial coding of the place cell population is a determinant of the behavioral performance of the agent in the closed-loop simulations, regardless of the specific value of other place cell parameters.

**Fig. 9.**
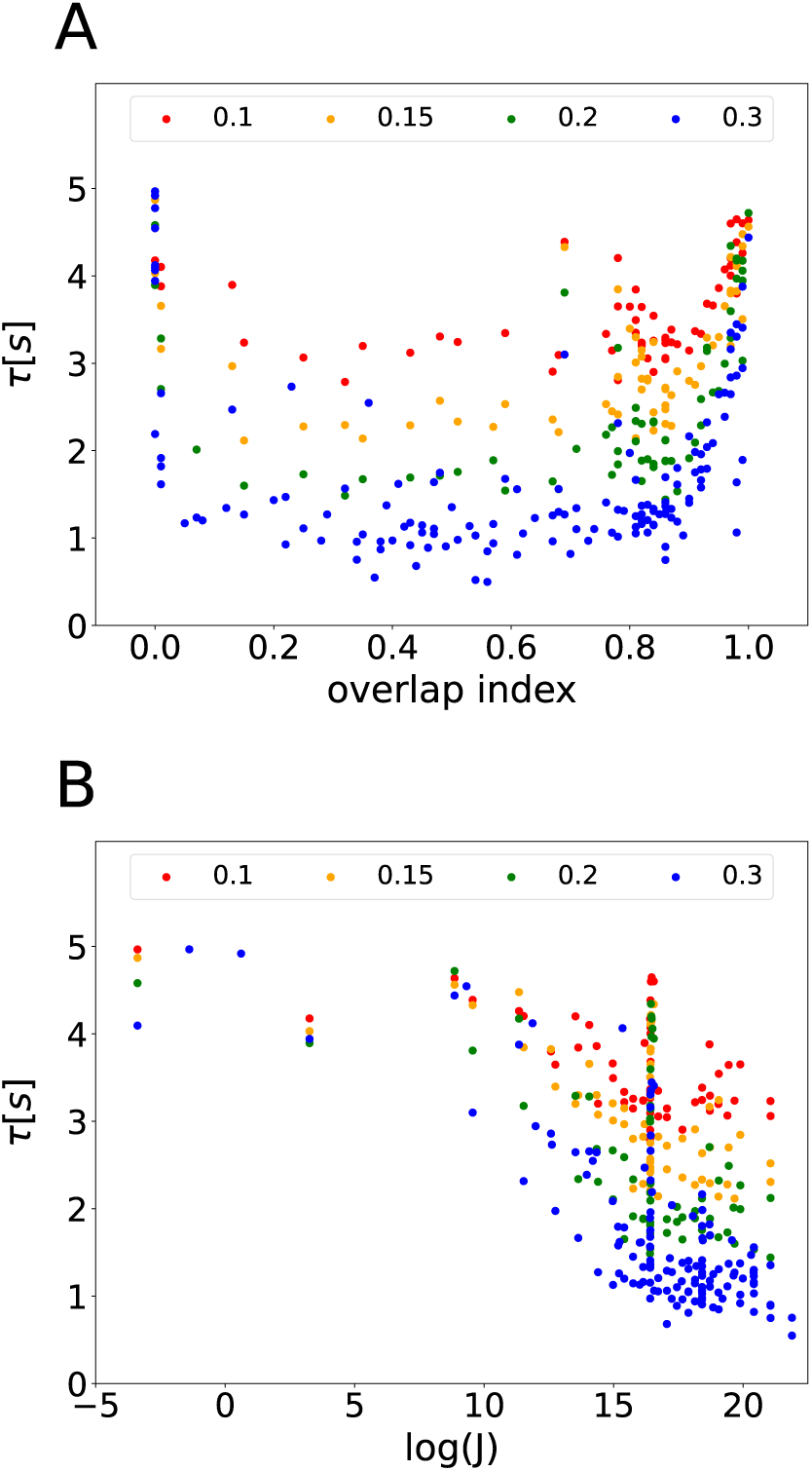
Summary of dependence of navigation performance on key parameters. Two variables are sufficient to account for task performance (as measured by escape latency) across all simulations: The reward zone radius (indicated by color) and spatial coding efficiency. **A**: The dependence of latency on reward zone size and overlap index is consistent across all simulations. The best performance is achieved around overlap index= 0.5, regardless of goal size. **B**: The dependence of latency on Fisher information is also consistent across all simulations, albeit somewhat more noisy. However, the relationship is monotonic and therefore the Fisher information captures a more fundamental property that is important for spatial navigation, i.e. the efficiency of spatial coding.

## 4 Discussion

Using closed-loop simulations of spatial learning based on spiking neural networks and reward-driven STDP, we studied how the agent’s navigation performance depends on the goal size and the parameters of the place cell population. While the dependence on goal size is straight forward, the larger the better, the dependence on cell number and field size is complex and difficult to interpret. We found that the Fisher information of the place cell population, which measures the efficiency of spatial coding, provided a simple correlate of navigation performance. Therefore, our work not only shows that the efficient coding hypothesis can be applied fruitfully to spatial representations, it also demonstrates that efficient spatial coding is important for function, i.e. spatial navigation and learning, as well.

### 4.1 Related Closed-Loop Simulation Work

The closest simulation framework to ours is the one that our model was based on Brzosko et al. (2017). That study used the model to study the role of sequential modulation of synaptic plasticity rule in learning a navigation task, but did not study the role of place cell parameters. Another closed-loop simulator based on spiking neural networks, SPORE Kaiser et al. (2019), provides an interface between NEST (Gewaltig & Diesmann, 2007) and the GAZEBO robotics simulator^1^, but primarily targets robotics tasks. Furthermore, there is a framework for simulating closed-loop behavior based on reinforcement learning that is oriented towards neuroscience (CoBeL-RL) (Walther et al., 2021). While this allows for studying the computational solutions that emerge under different constraints, such as place-cell like representations Vijayabaskaran and Cheng (2022), how the statistics of hippocampal replay emerges in the network Diekmann and Cheng (2022), or the function of memory replay Zeng, Wiskott, and Cheng (2022), these studies were based on machine learning implementations of reinforcement learning and neural networks. Even though they help us understand the abstract computations performed by the brain during spatial navigation, the internal processes of this model significantly differ from those in the brain. In addition, there are simulation frameworks that target machine learning or industrial applications, such as, for example, DeepMind Lab (Beattie et al., 2016), RLlib (Liang et al., 2018), MiniGrid^2^, and MAgents (Zheng et al., 2018).

### 4.2 Efficiency of Spatial Coding

Spatial coding by individual place cells has been quantified since the early 1990’s by the spatial information. It is based on Shannon information and was introduced to hippocampal research by Skaggs, Mcnaughton, and Gothard (1992). However, spatial information has only been used to quantify the degree of spatial modulation of one place cell’s activity map, but not the spatial coding efficiency in the population.

The more detailed results aside, our results imply a simpler relationship to roughly maintain a similar level of coding efficiency in a limited parameter range: the smaller the fields are, the larger the number of fields should be to maintain the same level of coding efficiency. This principle accords well with experimental observations and provides an explanation for the observation that when smaller place fields are observed in an area of an environment, the density of fields there is higher too (Ainge et al., 2007; Dupret et al., 2010; Gauthier & Tank, 2018; Grieves et al., 2018, 2016; Hollup et al., 2001; Jarzebowski et al., 2022; Kaufman et al., 2020; I. Lee et al., 2006; J.S. Lee et al., 2020; Sato et al., 2020; Tanni et al., 2022; Tryon et al., 2017; Turi et al., 2019; Zaremba et al., 2017).

### 4.3 Limits of Efficient Coding

However, the observation that field sizes vary systematically with environmental features and behavior (Parra-Barrero et al., 2021), suggests that absolute field sizes do matter. If the Fisher information was the only variable that mattered for navigation performance, the hippocampus could as well have allocated a different number of fields with different sizes as long as the Fisher information is preserved. However, the brain consistently opts for more and smaller fields in certain situations, so this suggests that there are factors beyond the Fisher information that constrain spatial coding in the place cell population. Some variables that might impose additional constraints are the size of obstacles (our simulation did not include any), and temporal coding linked to the behavior of the animal, such as running speed (Parra-Barrero et al., 2021). In addition, there are more generic limitations on efficient coding, i.e. energy consumption (Crotty, Lasker, & Cheng, 2012), and limits on peak firing rate of neurons due to spike dynamics. Future work is required to explore these factors and the trade-off between them.

4.4 Learning vs. Navigation

The current study distinguishes between spatial navigation and spatial learning. Here, we define spatial navigation as the set of movements that an agent makes to reach certain locations in an environment. Spatial learning is the acquisition of spatial information and/or the process of determining which movements are most beneficial for the agent. In our model, the agent does the latter, i.e. it learns which actions to perform in certain spatial locations. This corresponds to what has been called a response strategy. In our model, there are limitations on spatial navigation because of the architecture of the neural network and the inputs it receives (from place cells). For instance, due to the finite number of action selection neurons and the exponential kernel in the output (Eq. 6), the agent’s movements have a limited temporal and spatial precision, i.e. they cannot be arbitrarily precise. Furthermore, there are limitations on spatial learning. Even if the network weights could, in principle, be set up such that it would perform optimal navigation, the synaptic learning rule and the statistics of learning might not be able to bring about such network weights. In other words, a solution might exist, but the learning process cannot find this rule.

To begin dissociating the limitations on learning from those on navigation itself, we compared the performance of trained networks to randomly initialized networks. We found that some difference in performance was already present in the random network before learning commenced (Fig. A4), suggesting that for some place cell parameters, navigation is inherently more robust than for others and that learning amplifies this difference. However, it remains an open question whether other ways of optimizing synaptic weights, i.e., avoiding learning via reward-driven STDP, might yield a better solution for spatial navigation. In cases, where this is possible, the limitation of the network is primarily on learning, not navigation. In cases, where a well-performing solution cannot be found by other means either, the limitation is on navigation *per se*. This requires a more systematic exploration of different optimization algorithms that is beyond the scope of this paper.

### 4.5 Conclusion

While the encoding of sensory information has received a great deal of attention in neuroscience, the functional role of sensory representations is much less clear. This applies especially to spatial representations and navigation. Here, we have shown using computational modeling that the efficiency of spatial coding in the inputs directly influences the navigation performance of an agent based on spiking neural networks and reward-modulated STDP.

## Declarations

### Code availability

The CoBeL-spike framework is available at https://github.com/sencheng/CoBeL-spike.

### Authors’ contributions

All authors contributed to conception and design of the framework and study. Mohammadreza Mohagheghi Nejad wrote the code. Behnam Ghazinouri performed and analyzed simulations. Behnam Ghazinouri and Sen Cheng wrote the first draft of the manuscript. All authors edited and approved the submitted version.

## Acknowledgements

We thank Amit Pal, Kolja Dorschel, Paul Adler, Raymond Black and Yorick Sens for help with the figures and the CoBeL-spike software framework.

## Appendix A Supplementary materials

**Fig. A1.**
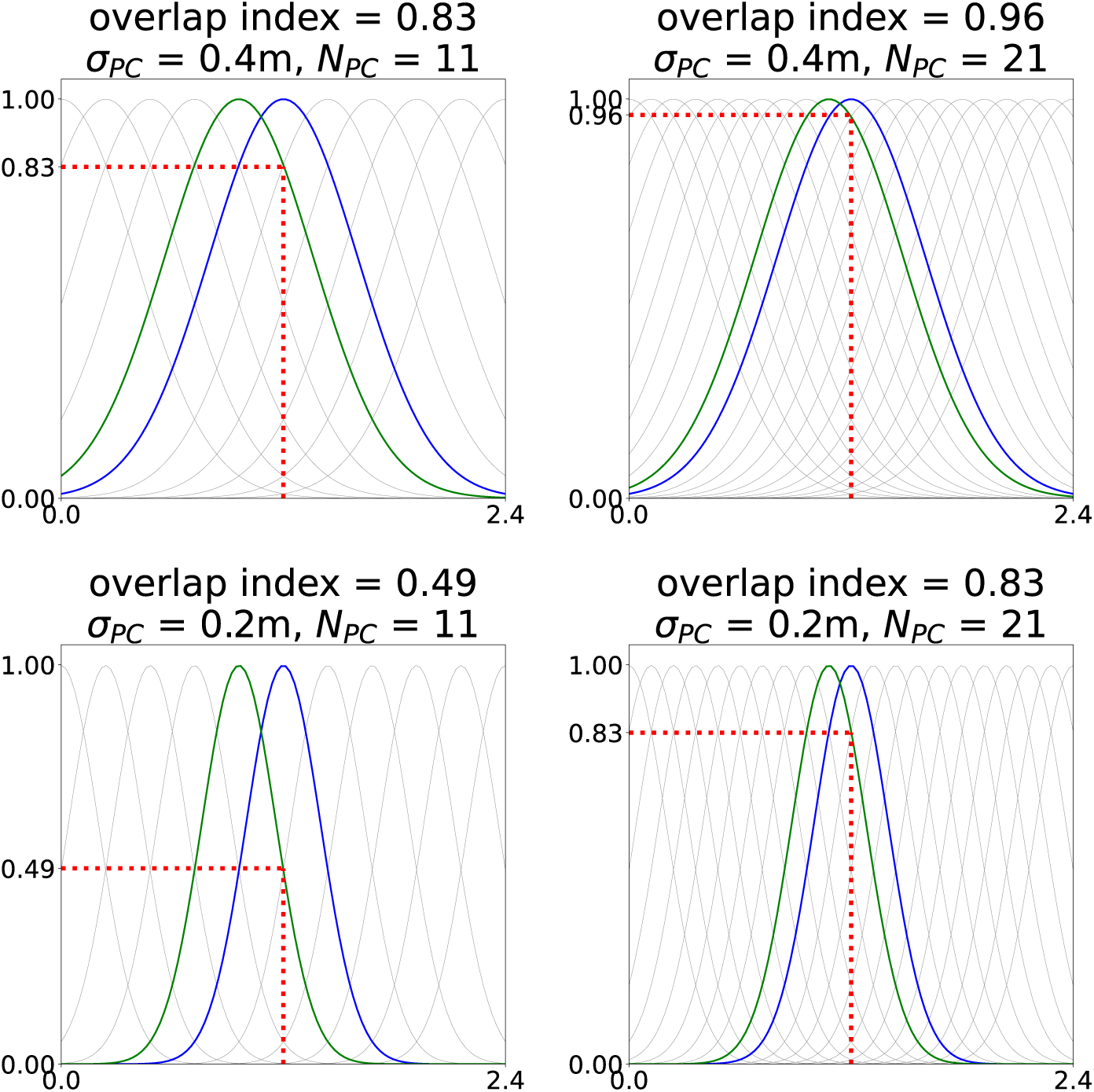
Illustration of the overlap index in 1-d. Bottom-Left: tuning curves of *N_PC_* = 11 place cells with field size *σ_PC_* = 0.2, homogeneously distributed over one edge of the environment (2.4 m). The overlap index is defined as the relative firing rate of the nearest neighboring place cell (green) in the center of a given place cell (blue). In this example, overlap index= 0.49. The overlap index increases with the cell number (left to right) and with the place field size (bottom to top). Since both cases scale in the same way (overlap index= 0.83), one can maintain a constant overlap, if decreasing the cell number and increasing field size (or vice a versa) by the same factor, i.e., such that *N_PC_ σ_PC_* = *const*.

**Fig. A2.**
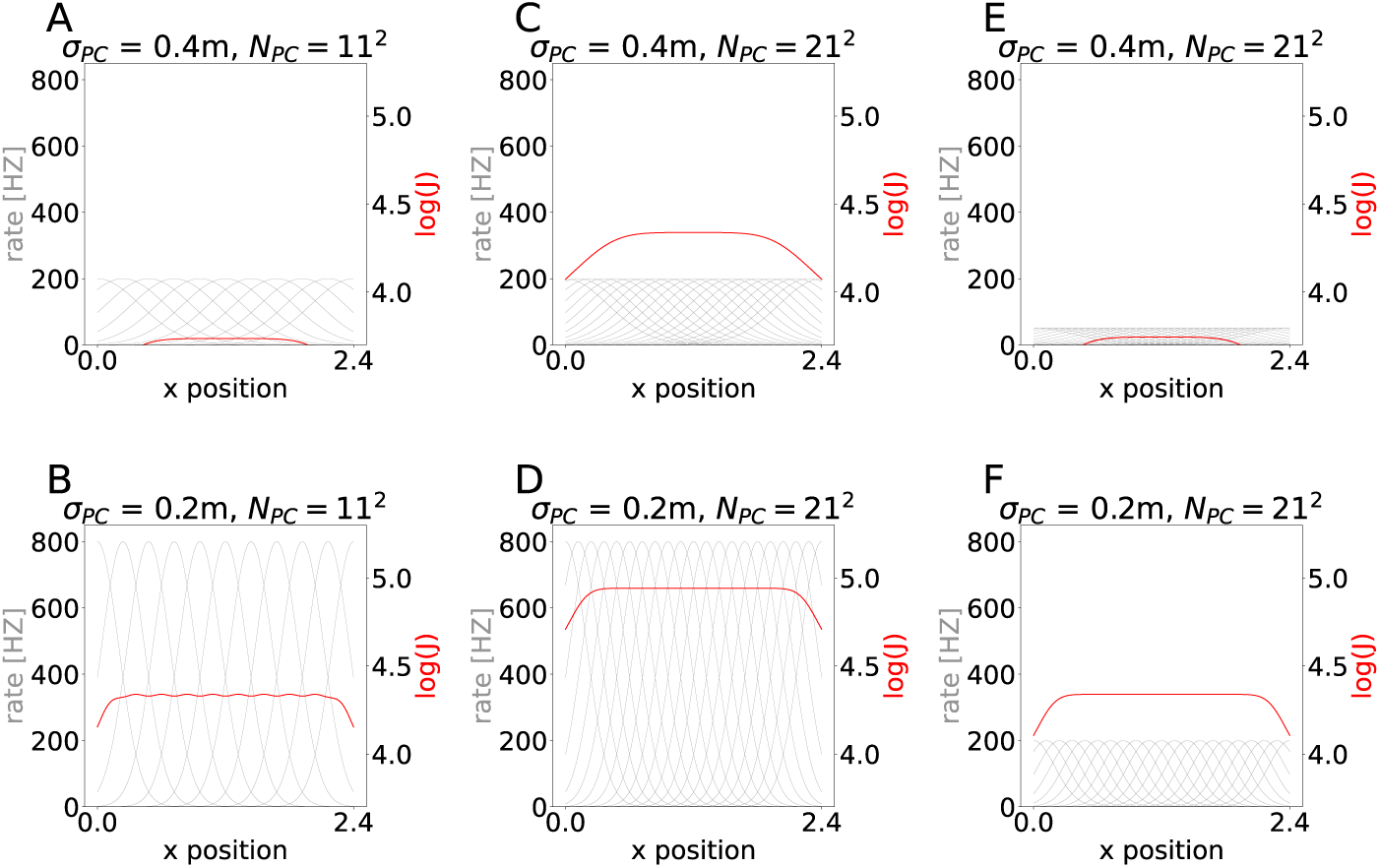
Illustration of Fisher information. Binary logarithm of Fisher information for six place cell populations characterized by different values of cell number, field size, and maximum individual firing rate. Generally, larger field sizes leads to lower Fisher information (comparing top row to bottom row) and increasing the number of cells increases the Fisher information (comparing 2nd column to 1st). However, in our simulations we keep the summed network activity in the center roughly constant (3500 Hz) by decreasing the individual maximum firing rate *η_PC_*. This scaling reduces the Fisher information (comparing 3rd column to 2nd). The effects of increasing cell number and decreasing maximum firing rate balance each other (comparing 1st column to 3rd). As a result, scaling cell number in our simulations does not influence the Fisher information under normal circumstances.

**Fig. A3.**
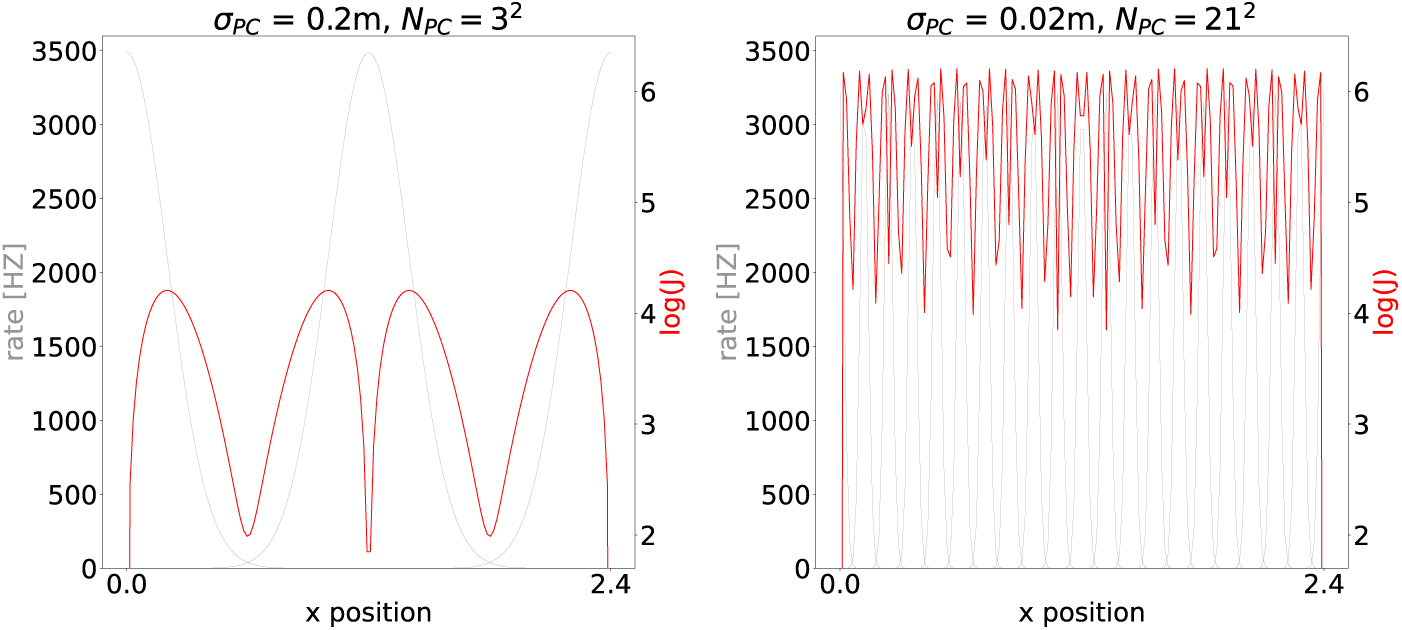
Fisher information fluctuates greatly when place fields are sparse. The Fisher information for most values of the place cell parameters is fairly homogeneous across the environment. However, that is not the case for very low cell numbers or very small field sizes. Therefore, we chose the minimum Fisher information along the path from the start point to the goal’s center to quantify spatial coding by the place cell population, because the locations where Fisher information is low are the bottlenecks of spatial learning. This figure illustrates two examples of extremely heterogeneous Fisher information.

**Fig. A4.**
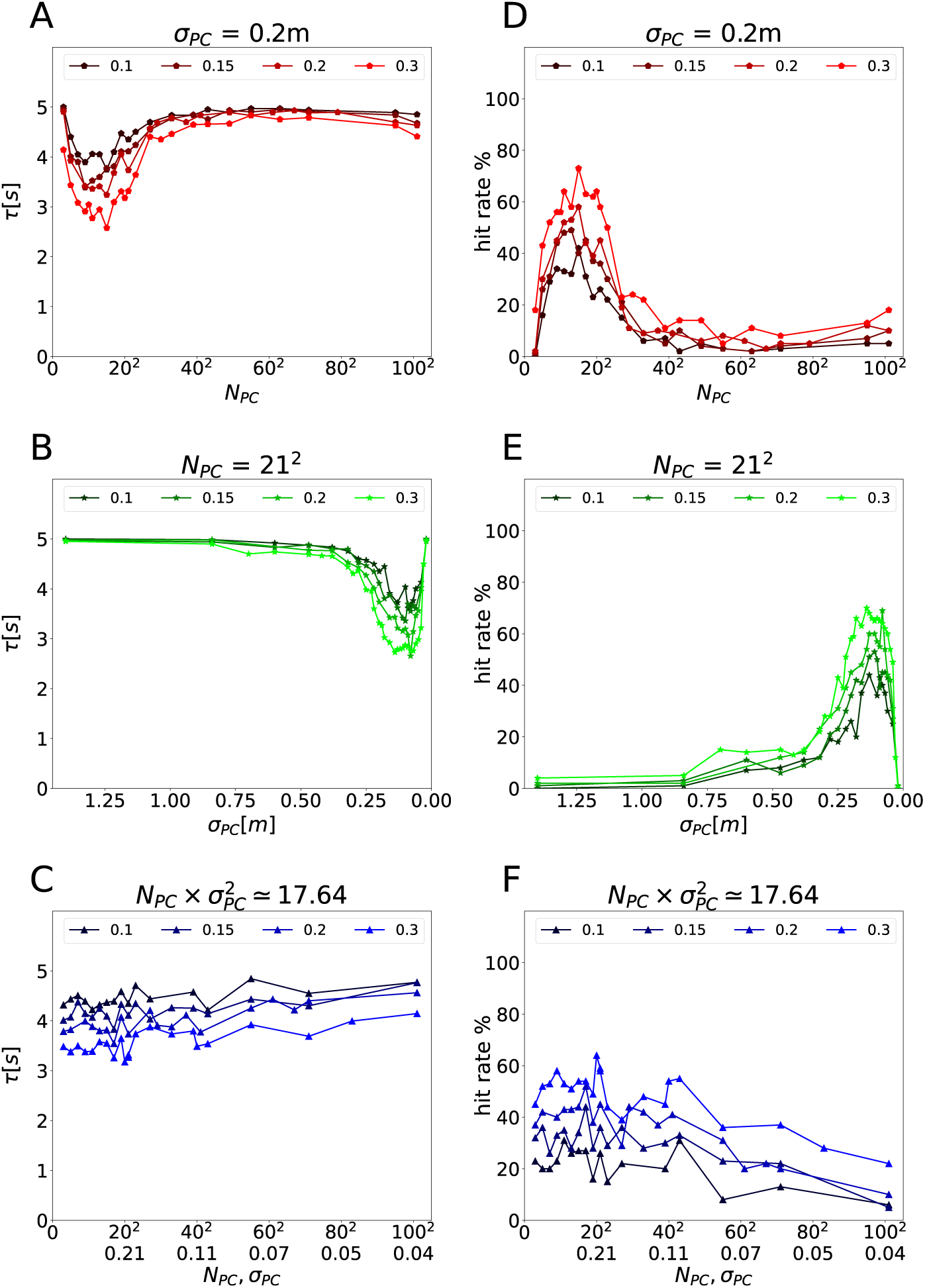
Navigation performance before learning based on randomly initialized network. Escape latency as a function **A**: of place cell number *N_PC_* for fixed *σ_PC_* = 0.2 m, **B**: of place field size *σ_PC_* for fixed *N_PC_* = 21^2^, and **C**: of both variables scaled simultaneous such that 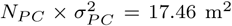. Each point represents the average trial duration over 100 repetitions. The color represents different goal radii (as indicated). **D-F**: The hit rate, i.e. the fraction of trials in which the agent successfully navigates to the goal, shown using the same plotting convention as in A-C.

## Footnotes

1 http://gazebosim.org/

2 https://github.com/Farama-Foundation/Minigrid

## References

1. Abbott, L.F., & Dayan, P. (1999). The effect of correlated variability on the accuracy of a population code. Neural computation, 11 (1), 91–101.

2. Ainge, J.A., Tamosiunaite, M., Woergoetter, F., Dudchenko, P.A. (2007). Hippocampal ca1 place cells encode intended destination on a maze with multiple choice points. Journal of Neuroscience, 27 (36), 9769–9779.

3. Barlow, H.B. (2001). Redundancy reduction revisited. Network: computation in neural systems, 12 (3), 241.

4. Barlow, H.B., et al. (1961). Possible principles underlying the transformation of sensory messages. Sensory communication, 1 (01).

5. Beattie, C., Leibo, J.Z., Teplyashin, D., Ward, T., Wainwright, M., Küttler, H., … others (2016). Deepmind lab. arXiv preprint arXiv:1612.03801.

6. Bernard, C., Ge, Y., Stockley, E., Willis, J., Wheal, H.V. (1994). Synaptic integration of nmda and non-nmda receptors in large neuronal network models solved by means of differential equations. Biological cybernetics, 70 (3), 267–273.

7. Brockman, G., Cheung, V., Pettersson, L., Schneider, J., Schulman, J., Tang, J., Zaremba, W. (2016). Openai gym. arXiv preprint arXiv:1606.01540.

8. Brunel, N., & Nadal, J.-P. (1998). Mutual information, fisher information, and population coding. Neural computation, 10 (7), 1731–1757.

9. Brzosko, Z., Zannone, S., Schultz, W., Clopath, C., Paulsen, O. (2017, July). Sequential neuromodulation of Hebbian plasticity offers mechanism for effective reward-based navigation. eLife, 6, e27756. 10.7554/eLife.27756

10. Crotty, P., Lasker, E., Cheng, S. (2012). Constraints on the synchronization of entorhinal cortex stellate cells. Physical Review E, 86 (1), 011908.

11. Diekmann, N., & Cheng, S. (2022). A model of hippocampal replay driven by experience and environmental structure facilitates spatial learning. bioRxiv.

12. Diesmann, M., Gewaltig, M.-O., Rotter, S., Aertsen, A. (2001). State space analysis of synchronous spiking in cortical neural networks. Neurocomputing, 38, 565–571.

13. Djurfeldt, M., Hjorth, J., Eppler, J.M., Dudani, N., Helias, M., Potjans, T.C., … Ekeberg, Ö (2010). Run-time interoperability between neuronal network simulators based on the music framework. Neuroinformatics, 8, 43–60.

14. Dupret, D., O’neill, J., Pleydell-Bouverie, B., Csicsvari, J. (2010). The reorganization and reactivation of hippocampal maps predict spatial memory performance. Nature neuroscience, 13 (8), 995–1002.

15. Eliav, T., Maimon, S.R., Aljadeff, J., Tsodyks, M., Ginosar, G., Las, L., Ulanovsky, N. (2021). Multiscale representation of very large environments in the hippocampus of flying bats. Science, 372 (6545), eabg4020.

16. Finkelstein, A., Ulanovsky, N., Tsodyks, M., Aljadeff, J. (2018). Optimal dynamic coding by mixed-dimensionality neurons in the head-direction system of bats. Nature communications, 9 (1), 1–17.

17. Fisher, R.A. (1922). On the mathematical foundations of theoretical statistics. Philosophical transactions of the Royal Society of London. Series A, containing papers of a mathematical or physical character, 222 (594-604), 309–368.

18. Florian, R.V. (2007). Reinforcement learning through modulation of spiketiming-dependent synaptic plasticity. Neural computation, 19 (6), 1468–1502.

19. Gauthier, J.L., & Tank, D.W. (2018). A dedicated population for reward coding in the hippocampus. Neuron, 99 (1), 179–193.

20. Gewaltig, M.-O., & Diesmann, M. (2007). Nest (neural simulation tool). Scholarpedia, 2 (4), 1430.

21. Graham, D.J., & Field, D.J. (2007). Efficient neural coding of natural images. New Encycl. Neurosci, 1, 1–18.

22. Grieves, R.M., Duvelle, É., Dudchenko, P.A. (2018). A boundary vector cell model of place field repetition. Spatial Cognition & Computation, 18 (3), 217–256.

23. Grieves, R.M., Wood, E.R., Dudchenko, P.A. (2016). Place cells on a maze encode routes rather than destinations. Elife, 5.

24. Hartley, T., Burgess, N., Lever, C., Cacucci, F., O’keefe, J. (2000). Modeling place fields in terms of the cortical inputs to the hippocampus. Hippocampus, 10 (4), 369–379.

25. Herzog, L.E., Pascual, L.M., Scott, S.J., Mathieson, E.R., Katz, D.B., Jadhav, S.P. (2019). Interaction of taste and place coding in the hippocampus. Journal of Neuroscience, 39 (16), 3057–3069.

26. Hintjens, P. (2013). Zeromq: messaging for many applications. “O’Reilly Media, Inc.”.

27. Hollup, S.A., Molden, S., Donnett, J.G., Moser, M.-B., Moser, E.I. (2001). Accumulation of hippocampal place fields at the goal location in an annular watermaze task. Journal of Neuroscience, 21 (5), 1635–1644.

28. Izhikevich, E.M. (2007). Solving the distal reward problem through linkage of stdp and dopamine signaling. Cerebral cortex, 17 (10), 2443–2452.

29. Jack, J., Noble, D., Tsien, R. (1983). Electrical current flow in excitable cells oxford university press. Oxford University Press, Oxford.

30. Jarzebowski, P., Hay, Y.A., Grewe, B.F., Paulsen, O. (2022). Different encoding of reward location in dorsal and intermediate hippocampus. Current Biology, 32 (4), 834–841.

31. Jordan, J., Weidel, P., Morrison, A. (2019). A Closed-Loop Toolchain for Neural Network Simulations of Learning Autonomous Agents. Frontiers in Computational Neuroscience, 13, 46. 10.3389/fncom.2019.00046

32. Kaiser, J., Hoff, M., Konle, A., Vasquez Tieck, J.C., Kappel, D., Reichard, D., … others (2019). Embodied synaptic plasticity with online reinforcement learning. Frontiers in Neurorobotics, 13, 81.

33. Kaufman, A.M., Geiller, T., Losonczy, A. (2020). A role for the locus coeruleus in hippocampal ca1 place cell reorganization during spatial reward learning. Neuron, 105 (6), 1018–1026.

34. Kjelstrup, K.B., Solstad, T., Brun, V.H., Hafting, T., Leutgeb, S., Witter, M.P., … Moser, M.-B. (2008). Finite scale of spatial representation in the hippocampus. Science, 321 (5885), 140–143.

35. Kloosterman, F., Layton, S.P., Chen, Z., Wilson, M.A. (2014). Bayesian decoding using unsorted spikes in the rat hippocampus. Journal of neurophysiology.

36. Kobayashi, R., Tsubo, Y., Shinomoto, S. (2009). Made-to-order spiking neuron model equipped with a multi-timescale adaptive threshold. Frontiers in computational neuroscience, 3, 9.

37. Lee, I., Griffin, A.L., Zilli, E.A., Eichenbaum, H., Hasselmo, M.E. (2006). Gradual translocation of spatial correlates of neuronal firing in the hippocampus toward prospective reward locations. Neuron, 51 (5), 639–650.

38. Lee, J.S., Briguglio, J.J., Cohen, J.D., Romani, S., Lee, A.K. (2020). The statistical structure of the hippocampal code for space as a function of time, context, and value. Cell, 183 (3), 620–635.

39. Legenstein, R., Pecevski, D., Maass, W. (2008). A learning theory for reward-modulated spike-timing-dependent plasticity with application to biofeedback. PLoS computational biology, 4 (10), e1000180.

40. Liang, E., Liaw, R., Nishihara, R., Moritz, P., Fox, R., Goldberg, K., … Stoica, I. (2018). Rllib: Abstractions for distributed reinforcement learning. International conference on machine learning (pp. 3053–3062).

41. Mathis, A., Herz, A.V., Stemmler, M. (2012). Optimal population codes for space: grid cells outperform place cells. Neural computation, 24 (9), 2280–2317.

42. McNaughton, B.L., Battaglia, F.P., Jensen, O., Moser, E.I., Moser, M.-B. (2006). Path integration and the neural basis of the’cognitive map’. Nature Reviews Neuroscience, 7 (8), 663–678.

43. Morris, R.G. (1981). Spatial localization does not require the presence of local cues. Learning and motivation, 12 (2), 239–260.

44. O’Keefe, J., & Nadel, L. (1978). The hippocampus as a cognitive map. Clarendon Press.

45. Parra-Barrero, E., Diba, K., Cheng, S. (2021). Neuronal sequences during theta rely on behavior-dependent spatial maps. Elife, 10, e70296.

46. Potjans, W., Morrison, A., Diesmann, M. (2010). Enabling functional neural circuit simulations with distributed computing of neuromodulated plasticity. Frontiers in computational neuroscience, 4, 141.

47. Ralf, H., & Bethge, M. (2010). Evaluating neuronal codes for inference using fisher information. Advances in neural information processing systems, 23.

48. Rich, P.D., Liaw, H.-P., Lee, A.K. (2014). Large environments reveal the statistical structure governing hippocampal representations. Science, 345 (6198), 814–817.

49. Rieke, F., Warland, D., Van Steveninck, R.d.R., Bialek, W. (1999). Spikes: exploring the neural code. MIT press.

50. Rolls, E.T., & Treves, A. (2011). The neuronal encoding of information in the brain. Progress in neurobiology, 95 (3), 448–490.

51. Rotter, S., & Diesmann, M. (1999). Exact digital simulation of time-invariant linear systems with applications to neuronal modeling. Biological cybernetics, 81 (5), 381–402.

52. Samsonovich, A., & McNaughton, B.L. (1997). Path integration and cognitive mapping in a continuous attractor neural network model. Journal of Neuroscience, 17 (15), 5900–5920.

53. Sato, M., Mizuta, K., Islam, T., Kawano, M., Sekine, Y., Takekawa, T., … others (2020). Distinct mechanisms of over-representation of landmarks and rewards in the hippocampus. Cell reports, 32 (1), 107864.

54. Simoncelli, E.P. (2003). Vision and the statistics of the visual environment. Current opinion in neurobiology, 13 (2), 144–149.

55. Skaggs, W., Mcnaughton, B., Gothard, K. (1992). An information-theoretic approach to deciphering the hippocampal code. Advances in neural information processing systems, 5.

56. Spreizer, S., Aertsen, A., Kumar, A. (2019). From space to time: Spatial inhomogeneities lead to the emergence of spatiotemporal sequences in spiking neuronal networks. PLoS computational biology, 15 (10), e1007432.

57. Tanni, S., De Cothi, W., Barry, C. (2022). State transitions in the statistically stable place cell population correspond to rate of perceptual change. Current Biology, 32 (16), 3505–3514.

58. Taube, J.S. (1998). Head direction cells and the neurophysiological basis for a sense of direction. Progress in neurobiology, 55 (3), 225–256.

59. Tryon, V.L., Penner, M.R., Heide, S.W., King, H.O., Larkin, J., Mizumori, S.J. (2017). Hippocampal neural activity reflects the economy of choices during goal-directed navigation. Hippocampus, 27 (7), 743–758.

60. Turi, G.F., Li, W.-K., Chavlis, S., Pandi, I., O’Hare, J., Priestley, J.B., … others (2019). Vasoactive intestinal polypeptide-expressing interneurons in the hippocampus support goal-oriented spatial learning. Neuron, 101 (6), 1150–1165.

61. van Wijngaarden, J.B., Babl, S.S., Ito, H.T. (2020). Entorhinal-retrosplenial circuits for allocentric-egocentric transformation of boundary coding. Elife, 9, e59816.

62. Vijayabaskaran, S., & Cheng, S. (2022). Navigation task and action space drive the emergence of egocentric and allocentric spatial representations. PLOS Computational Biology, 18 (10), e1010320.

63. Walther, T., Diekmann, N., Vijayabaskaran, S., Donoso, J.R., Manahan-Vaughan, D., Wiskott, L., Cheng, S. (2021). Context-dependent extinction learning emerging from raw sensory inputs: A reinforcement learning approach. Scientific Reports, 11 (1), 1–14.

64. Wiskott, L., & Sejnowski, T.J. (2002). Slow feature analysis: Unsupervised learning of invariances. Neural computation, 14 (4), 715–770.

65. Yamauchi, S., Kim, H., Shinomoto, S. (2011). Elemental spiking neuron model for reproducing diverse firing patterns and predicting precise firing times. Frontiers in computational neuroscience, 5, 42.

66. Zaremba, J.D., Diamantopoulou, A., Danielson, N.B., Grosmark, A.D., Kaifosh, P.W., Bowler, J.C., … Losonczy, A. (2017). Impaired hippocampal place cell dynamics in a mouse model of the 22q11. 2 deletion. Nature neuroscience, 20 (11), 1612–1623.

67. Zeng, X., Wiskott, L., Cheng, S. (2022). The functional role of episodic memory in spatial learning. bioRxiv, 2021–11.

68. Zheng, L., Yang, J., Cai, H., Zhou, M., Zhang, W., Wang, J., Yu, Y. (2018). Magent: A many-agent reinforcement learning platform for artificial collective intelligence. Proceedings of the aaai conference on artificial intelligence (Vol. 32).

